# Distinct subtypes of grey matter heterotopia show subtype-specific morpho-electric neuronal properties and dynamics of epileptiform activity in mice

**DOI:** 10.1101/2023.06.06.543853

**Authors:** Jean-Christophe Vermoyal, Delphine Hardy, Lucas Goirand-Lopez, Antonin Vinck, Lucas Silvagnoli, Aurélien Fortoul, Fiona Francis, Silvia Cappello, Ingrid Bureau, Alfonso Represa, Carlos Cardoso, Françoise Watrin, Thomas Marissal, Jean-Bernard Manent

## Abstract

Grey matter heterotopia (GMH) are neurodevelopmental disorders associated with abnormal cortical function and epilepsy. Subcortical band heterotopia (SBH) and periventricular nodular heterotopia (PVNH) are two well-recognized GMH subtypes in which neurons are misplaced, either forming nodules lining the ventricles in PVNH, or forming bands in the white matter in SBH. Although both PVNH and SBH are commonly associated with epilepsy, it is unclear whether these two GMH subtypes differ in terms of pathological consequences or, on the contrary, share common altered mechanisms. Here, we studied two robust preclinical models of SBH and PVNH, and performed a systematic comparative assessment of the physiological and morphological diversity of heterotopia neurons, as well as the dynamics of epileptiform activity and input connectivity. We uncovered a complex set of altered properties, including both common and distinct physiological and morphological features across heterotopia subtypes, and associated with specific dynamics of epileptiform activity. Taken together, these results suggest that pro-epileptic circuits in GMH are, at least in part, composed of neurons with distinct, subtype-specific, physiological and morphological properties depending on the heterotopia subtype. Our work supports the notion that GMH represent a complex set of disorders, associating both shared and diverging pathological consequences, and contributing to forming epileptogenic networks with specific properties. A deeper understanding of these properties may help to refine current GMH classification schemes by identifying morpho-electric signatures of GMH subtypes, to potentially inform new treatment strategies.

## Introduction

Disrupted formation of the cerebral cortex results in malformations of cortical development (MCDs), a large group of neurodevelopmental disorders associated with intellectual disability and epilepsy.^1^ Fueled with the rapid advances in neuroimaging and molecular genetics, around 200 MCD subtypes are now recognized, and classified according to the presumably altered developmental mechanisms, associated causative genes and imaging features. Hence, MCDs are sorted into 3 main groups based on altered neurodevelopmental steps, as malformations secondary to abnormal proliferation and apoptosis; neuronal migration disorders; and malformations of post-migrational cortical organization and connectivity.^2–4^

Grey matter heterotopia (GMH) belongs to the group of neuronal migration disorders, and histologically refers to masses of neurons found in an abnormal location and unable to migrate to their normal position in the cortex. Two well-recognized GMH subtypes are periventricular nodular heterotopia (PVNH) and subcortical band heterotopia (SBH). PVNH describes nodular masses of neurons located below the white matter along the ventricular walls, and protruding into the ventricles. SBH describes a band of aggregated neurons separated from the overlying cortex and lateral ventricles by layers of white matter.^3,4^ Aside from these well-described and classified GMH subtypes, a third group of very rare forms of subcortical heterotopia (SUBH) comprises heterotopia located within the white matter, between the cortex and lateral ventricles, but anatomically distinct from SBH.^5,6^ These rare forms of SUBH are however not considered as neuronal migration disorders as they are thought to result from abnormal proliferation of neurons and glia, or from abnormal post-migrational development.^5^

Although GMH are widely associated with altered cortical function and epilepsy,^1^ their pathophysiological bases remain poorly understood. This lack of knowledge regarding the morpho-functional properties and circuit organization of heterotopia neurons is largely due to a limited access to human tissue. Several rodent models with either a PVNH-like or a SBH-like phenotype have been described.^7–20^ However, no comparative characterization of the structural and functional properties of heterotopia neurons has been carried out so far in these two main GMH subtypes. In addition to increasing knowledge of the underlying mechanisms leading to epilepsy, such a comparative assessment could help to clarify whether some altered properties are specific to one GMH subtype, while other properties may be common to all GMH. This increased knowledge would potentially improve classification schemes and inform new treatment strategies or tailored therapies.^21^

To address this issue, we studied two GMH mouse models harboring PVNH-like and SBH-like malformations. We selected the Eml1^10^ and the RhoA^9^ conditional knockout (cKO) mice as they faithfully recapitulate these GMH phenotypes. The Eml1 gene encodes a microtubule-associated protein, and the RhoA gene encodes a small GTPase, and, interestingly, their deletion in mice leads to abnormal morphologies and distributions of radial glia progenitors, and results in impaired neuronal migration and heterotopia formation.^9,22^ However, the anatomical characteristics of the resulting heterotopia are strikingly different in the two models, one showing a SBH-like phenotype and the other a PVNH-like phenotype. To compare the structural and functional properties of heterotopia neurons in the two models, we performed a series of electrophysiological experiments aimed at assessing their physiological properties at both the cellular and network levels, and combined them with neuroanatomical methods. We uncovered a complex set of altered properties, including both common and distinct physiological and morphological features across heterotopia subtypes, and associated with specific dynamics of epileptiform activity. Taken together, these results suggest that pro-epileptic circuits in GMH are, at least in part, composed of neurons with distinct, subtype-specific, physiological and morphological properties depending on the heterotopia subtype.

## Material and methods

### Animals

Both GMH models are conditional knockout (cKO) mice obtained by crossing Emx1-Cre mouse lines^23,24^ with mice carrying loxP sites flanking exon 3 of the RhoA gene,^9,25^ and with mice carrying loxP sites flanking exon 8 of the Eml1 gene.^10^ All experiments were performed in agreement with European directive 2010/63/UE and received approval [2020080610441911_v2(APAFIS#26835)] from the French Ministry for Research after ethical evaluation by the Institutional Animal Care and Use Committee of Aix-Marseille University.

### Immunohistochemistry

Postnatal (P) 15 (P15) mice were deeply anesthetized with tiletamine/zolazepam (Zoletil, 40 mg/kg) and medetomidine (Domitor, 0.4 mg/kg). Depending on the antibody used for the immunohistochemistry, they were perfused transcardially using either Antigenfix (DiaPath) for anti-Ctip2, anti-Cux1 or anti-NeuN immunohistochemistry, or Antigenfix with 0.5% glutaraldehyde (Sigma-Aldrich) for anti-GABA immunohistochemistry. Brains were dissected out and post-fixed in Antigenfix overnight at 4°C. Serial frontal sections (100 μm) were performed using a vibrating microtome (Leica) and processed as free-floating sections for antigen-retrieval treatment and immunohistochemistry. For antigen-retrieval, sections were heated in 10 mM sodium citrate buffer (Ph6.0) with 0.05% Tween 20 for 30min at 100°C, and washed twice in phosphate-buffered saline (PBS) buffer at room temperature. For immunohistochemistry, sections were permeabilized in PBS with 0.3% Triton X-100 for 30 min at room temperature, blocked in PBS with 0.3% Triton X-100, 2% bovine serum albumin (BSA) and 3% normal goat serum for 1 h at room temperature, and incubated in PBS with 0.3% Triton X-100, 0.1% BSA and 1% normal goat serum overnight at 4°C with primary antibodies: rat anti-Ctip2 (1/500; Abcam, #ab18465), rabbit anti-Cux1 (1/500; SantaCruz, #sc-13024), rabbit anti-GABA (1/250; Sigma-Aldrich, #A2052), mouse anti-NeuN (1/1000; Millipore, #mab377). Sections were washed in PBS and incubated for 2 h at room temperature with secondary antibodies: goat anti-rat Alexa Fluor 555 (1/500; Invitrogen, #A21434), goat anti-rabbit Alexa Fluor 488 (1/500; Invitrogen, #A11034) and/or goat anti-mouse Alexa Fluor 555 (1/500; Invitrogen, #A31570). Sections were washed in PBS, counterstained with Hoechst 33342 (1/1000, Thermo Fisher Scientific) and mounted in Fluoromount (Thermo Fisher Scientific). Sections were imaged as mosaics on a wide-field upright microscope equipped with a structured illumination system (Apotome 2, Zeiss) and a 10X objective. Sections used for electrophysiology (see below) were fixed overnight in AntigenFix before biocytin was revealed with Alexa647-conjugated-streptavidin (1/200, Jackson ImmunoResearch). Images of biocytin-filled neurons were acquired as 3D stacks on a confocal microscope (Leica) with a ×40 oil objective.

### Image analysis

To analyze the cell type composition in the two heterotopia subtypes, regions of interest (ROI) corresponding to the heterotopia were extracted using Fiji 2.10 software^26^ from fluorescent images. Color channels were analyzed separately. Each ROI image was processed using the Noise2Void image denoising algorithm.^27^ Segmentation of immune-positive cells was performed using Cellpose 2.0.5.^28^ We used the human-in-the-loop training feature of Cellpose to improve the detection model on a training dataset of 30 images, as described.^29^ These training images were taken from random areas across all experimental conditions. Using this fine-tuned model, cells were segmented on all sections to quantify each cell type in the ROIs separately. To compare cell populations across different conditions, we calculated the number of cells per square millimeter. Tridimensional brain reconstructions were made using Free-D software^30^ from serial frontal sections taken with a wide-field upright microscope (Zeiss). Biocytin-filled neurons were reconstructed tree-dimensionally using Neurolucida 360 software (MBF Bioscience) from 3D stack images. Morphological parameters were extracted from neuronal reconstructions with L-Measure software.^31^

### Slice electrophysiology

Juvenile (P12-P15) mice of both sexes were deeply anesthetized with tiletamine/zolazepam (Zoletil, 40 mg/kg) and medetomidine (Domitor, 0.6 mg/kg), and decapitated. The brain was then quickly removed and was placed in chilled and oxygenated ACSF containing the following (in mM): 25 NaHCO3, 1,25 NaH2PO4.H2O, 6,3 D-glucose, 2.5 KCl, 7 MgCl2.6H2O, 0,5 CaCl2.2H2O and 132,5 choline chloride. Frontal slices (250-μm-thick) were obtained using a vibrating microtome (Leica). During the electrophysiological experiments, the slices were perfused with oxygenated normal ACSF at a rate of 2 ml/min containing the following (in mM): 126 NaCl, 26 NaHCO3, 1,2 NaH2PO4.H2O, 6,3 D-Glucose, 3,5 KCl, 1,3 MgCl2.6H2O, 2 CaCl2.2H2O. Recordings were amplified using a Multiclamp 700B amplifier (Molecular Devices) and digitized using a Digidata 1440A (Molecular Devices). Patch electrodes (ranging from 6 to 9 MΩ) were filled with intracellular solution containing (in mM): 120 KMeSO4, 10 KCl, 10 HEPES, 8 NaCl, 4 Mg-ATP, 0.3 Na-GTP, and 0,3 Tris-base (for recording intrinsic cell properties) or with 128 CsMeS, 10 HEPES, 10 Sodium PhosphoCreatine, 4 MgCl2, 4 NaATP, 0,4 NaGTP, 3 Ascorbic acid (for recording synaptic properties and laser scanning photo-stimulation, see below). The intracellular solution was also supplemented with 5 mM of biocytin to ensure laminar position and morphological reconstruction. The resting membrane potential (RMP) was recorded in current-clamp mode within the first 10 s after achieving whole-cell configuration without any current injection. The membrane resistance (Rm) and capacitance (Cm) were directly measured by Clampex (Molecular Devices) in voltage-clamp mode. The Ih current amplitudes were measured in voltage-clamp mode as the difference between the instantaneous current following each tested potential (from -130 mv to -60 mV) and the steady-state current at the end of each tested potential. The Ih current density was obtained by dividing the Ih current amplitude by Cm. The AP traces were obtained by injecting 500-ms long depolarizing currents steps from 0 to 500 pA in 10-pA increments. To be considered as an AP the depolarization should be higher than 0 mV. The AP threshold and peak amplitude were analyzed from the first AP induced by a 500-ms long minimal current. The AP threshold was defined as the membrane potential at which the first derivative of an evoked AP achieved 10% of its peak velocity (dV/dt). The AP peak amplitude was defined as the difference between the peak and baseline. The rheobase was defined as the minimum injected current required to trigger an AP.

### Calcium imaging

Brain of P12-P15 mice of both sexes were removed as described above and immersed in ice-cold, oxygenated, modified ACSF (in which 0.5 mM CaCl2 and 7 mM MgSO4; NaCl was replaced by an equimolar concentration of choline chloride). Next, frontal slices (380µm) were obtained using a Leica VT1200S vibratome. The loading of slices with the calcium indicator Fura2-AM was performed as described.^32,33^ In brief, slices were placed in a custom-made chamber,^34^ where they were incubated in 1 mL of normal oxygenated ACSF with 1mM Fura2-AM and 0.4% Pluronic F-127 in DMSO, and maintained for 30 minutes at 35-37°C in the dark. Then, slices were transferred for rest during 1 hour in oxygenated normal ACSF at room temperature (22°C). Prior to calcium imaging, slices are placed in a recording chamber and continuously perfused at a rate of 3ml/min with normal, oxygenated ACSF at 35-37°C. Calcium transient recording was performed using a multibeam multiphoton laser scanning system (LaVision Biotech, excitation wavelength 780nm) coupled to an Olympus microscope and a X10, NA-0.3 objective (Olympus). Images were acquired through a CCD camera (La Vision Imager 3QE) at 7,4 Hz (i.e., 136ms per frame with 2x2 binning and a pixel size of 600 nm). A typical imaging session lasted 7-15 minutes (i.e., 3000-6500 frames), and covered a field of 1720 x 1520 µm size situated on average 80µm below the surface (range: 50-100µm). We performed two movies per slice: a first in ACSF alone (a condition referred to as the baseline), then a second one after 5 minutes of treatment with 5µM of the GABAA receptor antagonist gabazine, to elicit epileptiform network activities. Cell detection and signal analysis was done using custom designed Matlab software (Math-Works, Natick, MA), as described in Marissal *et al.*^33^ In brief, we 1) semi-automatically detected the regions of interest (ROIs) matching the identified Fura-2-containing fluorescent neuronal somata within each subregion from each experimental condition, 2) semi-automatically identified each calcium event onsets and offsets, 3) calculated the proportion of active neurons, as well as the frequency of calcium transients, 4) detected synchronous Calcium Events (SCE), which were defined as activities during which the onset a significant fraction of neurons was activated within a short time window, 5) plotted time-correlation graphs that illustrated for each active neuron the mean time of activation (expressed as imaging frames) relative to all other cells and their cross-correlation scores. The zero-time reference corresponds to the peak of neuronal co-activation.

### Laser scanning photo-stimulation

Laser scanning photo-stimulation (LSPS) was performed as previously described in Bureau *et al.*^35^ and Plantier *et al.*^36^ Recirculating ACSF solution contained the following (in mM): 126 NaCl, 26 NaHCO3, 1,2 NaH2PO4.H2O, 6.3 D-Glucose, 3,5 KCl, 4 MgCl2, 4 CaCl2, 0,2 4-methoxy-7-nitroindolinyl (MNI)-caged glutamate (Tocris Bioscience), 0,005 CPP [()-3-(2-carboxypiperazin-4-yl)propyl-1-phosphonic acid] (NMDA receptors antagonist, Sigma Aldrich). Traces of whole-cell voltage-clamp recordings (holding potential, -70 mV) were sampled and filtered at 10 kHz. Photolysis of caged glutamate was performed with a 2-ms 20 mW pulse of a UV (355 nm) laser (DPSS Lasers Inc.) through a 0,16 NA 4 × objective (Olympus). The stimulus pattern for LSPS mapping consisted of 256 positions on a 16 × 16 grid (spacing = 75 μm). The laser moved in a spatial pattern designed to avoid consecutive glutamate uncaging over neighboring pixel sites. Synaptic input maps for individual neurons were constructed by computing the mean current amplitude calculated in a 100-ms time window 7-ms after the UV stimulus for each position of photostimulation. 3 maps were obtained per cell and averaged. Synaptic and direct responses were distinguished based on their latencies and direct responses (latency < 7ms) were excluded from the analysis.

### Statistics

The statistical analysis carried out is indicated in the legend of each figure. All tests were 2-tailed and the level of significance was set at p < 0.05. Hierarchical clustering and heatmaps of the morphological and physiological parameters were done with ClutVis as described by Metsalu and Vilo.^37^ Principal component analysis was performed with Prism 9 (GraphPad Software). Gardner-Altman estimation plots were generated as described in Ho *et al.*^38^ Bee swarm plots and rain cloud plots were generated using SuperPlotsOfData, as described in Goedhart.^39^ Violin SuperPlots were generated as described by Kenny and Schoen.^40^ Sample size was estimated using InVivoStat 4.7 based on power analysis and effect size.

### Data availability

The data that support the findings of this study are available from the corresponding author, upon reasonable request.

## Results

### General anatomy of the two grey matter heterotopia subtypes

We first characterized the heterotopia phenotypes in the two GMH mouse models using histological methods and tridimensional reconstructions. In the two models, we observed masses of neurons forming prominent grey matter heterotopia that extended bilaterally and rostrocaudally beneath the normally positioned (or normotopic) cortex. The general anatomy of heterotopia however varied between models. RhoA cKO mice harbored heterotopia that were located along the ventricular walls and protruded into the ventricles, especially at their most rostral ends, and were separated from the overlying cortex by the white matter (Fig 1A, left). Eml1 cKO mice harbored heterotopia that were located within the white matter, with layers of white matter separating them from both the overlying cortex and the ventricles (Fig 1A, right). Comparative morphometric measurements revealed that the overall heterotopia surface in Eml1 cKO mice is significantly larger than in RhoA cKO mice, especially at their most rostral positions (Fig 1B). In line with the neuropathological definitions and consensus classifications of MCDs and GMH,^3,4^ and in particular regarding the locations of heterotopia in relation to the white matter and ventricles, our observations suggest that heterotopia in RhoA cKO mice resemble human Periventricular Nodular Heterotopia (PVNH), and heterotopia in Eml1 cKO mice resemble human Subcortical Band Heterotopia (SBH).

**Figure 1.**
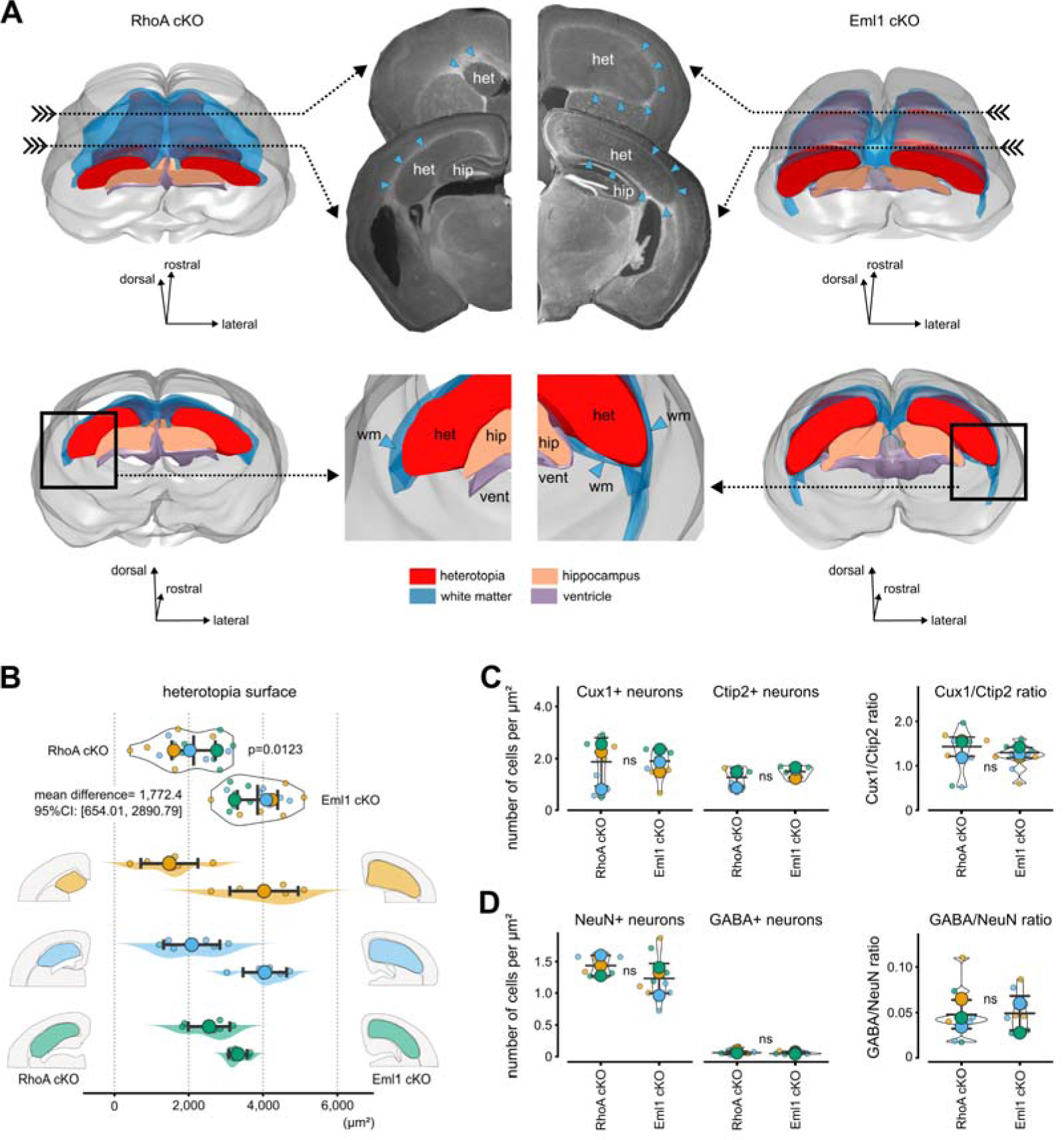
General anatomy of the two grey matter heterotopia subtypes. (**A**) Top views (top, left and right) and rear views (bottom, left and right) of the rostral part of threeldimensionally reconstructed brains from a RhoA cKO (left) and from a Eml1 cKO mouse (right). (middle, top) frontal sections taken at the indicated positions (arrows) showing the position of the heterotopia (het) relative to the white matter (blue arrowheads). (middle, bottom) enlarged views showing the position of the heterotopia relative to the white matter (wm, blue), hippocampus (hip, orange) and ventricles (vent, purple). **(B**) Bee swarm plots (top) of the heterotopia surface measured at 3 rostrocaudal levels (orange, blue and green) in RhoA (left) and Eml1 (right) cKO mice. Rain cloud plots (bottom) illustrating the data distribution at every rostrocaudal levels. (**C**) Bee swarm plots illustrating the density of Cux1+ and Ctip2+ neurons (left) and the ratio of Cux1/Ctip2 (right) in RhoA and Eml1 cKO mice. (**D**) Bee swarm plots illustrating the density of NeuN+ and GABA+ neurons (left) and the ratio of GABA/NeuN (right) in RhoA and Eml1 cKO mice. In B-D, small dots correspond to values of individual sections and large dots to the mean. Colors correspond to the same rostrocaudal levels as in B. Error bars show SD. Welch’s t-test, N=6 mice per GMH model.

### Both common and distinct aberrant neuronal morphologies are present in PVNH-like and SBH-like heterotopia

We next analyzed the cell type composition of the two heterotopia models following immunostainings with Cux1 and Ctip2 antibodies to estimate numbers of neurons with a superficial layer (L2-4) identity and those with a deep layer (L5-6) identity, respectively (Fig 1C). We also used antibodies against the neuron-specific nuclear protein NeuN and GABA, to estimate numbers of neurons and putative GABAergic interneurons, respectively (Fig 1D). We used the deep learning algorithm Cellpose to detect immunopositive cells and estimate their densities, and found that the two heterotopia subtypes were composed of roughly similar numbers of superficial and deep cortical neurons (Fig 1C), neurons and interneurons on average (Fig 1D).

To refine the analysis of the cell type composition in the two heterotopia subtypes, we filled heterotopia neurons with biocytin, and recovered their axodendritic morphologies after *post hoc* immunostainings and tridimensional reconstructions. To assess in an unbiased fashion whether similar or distinct morphological subtypes were present in the two heterotopia subtypes, we performed hierarchical clustering of the morphological parameters extracted from neuronal reconstructions (see material and methods and Fig 2A). We found that heterotopia neurons clustered into 3 morphological subtypes or ‘morphotypes’ (Fig 2B), two of them being selectively enriched in the heterotopia of Eml1 cKO mice (65% and 80%, respectively), and the third one mostly found in the heterotopia of RhoA cKO mice (75%). Principal component analysis (Fig 2C-H) confirmed these observations, and showed that while all morphotypes could be found across the 2 heterotopia subtypes, morphotypes 1 and 2 were selectively enriched in Eml1 cKO mice, while morphotype 3 was enriched in RhoA cKO mice. Overall, these unbiased methods revealed that grouping into morphotypes was influenced by the morphological features of distinct neuronal compartments. Hence, radar charts plotting the morphological parameters of axons, apical and basal dendrites for the 3 morphotypes (Fig 2I) showed that neurons of morphotype 1 had greater scores in axon-related features; that neurons of morphotype 2 had greater scores in parameters combining both apical and basal dendrites-related features; and that neurons of morphotype 3 had greater scores in parameters combining both apical dendritic and axonal features. A visual comparison of the reconstructed neuronal morphologies (examples are shown in Fig 2J) was consistent with this. Collectively, this suggests that both common and distinct aberrant neuronal morphologies are present in PVNH-like and SBH-like heterotopia.

**Figure 2.**
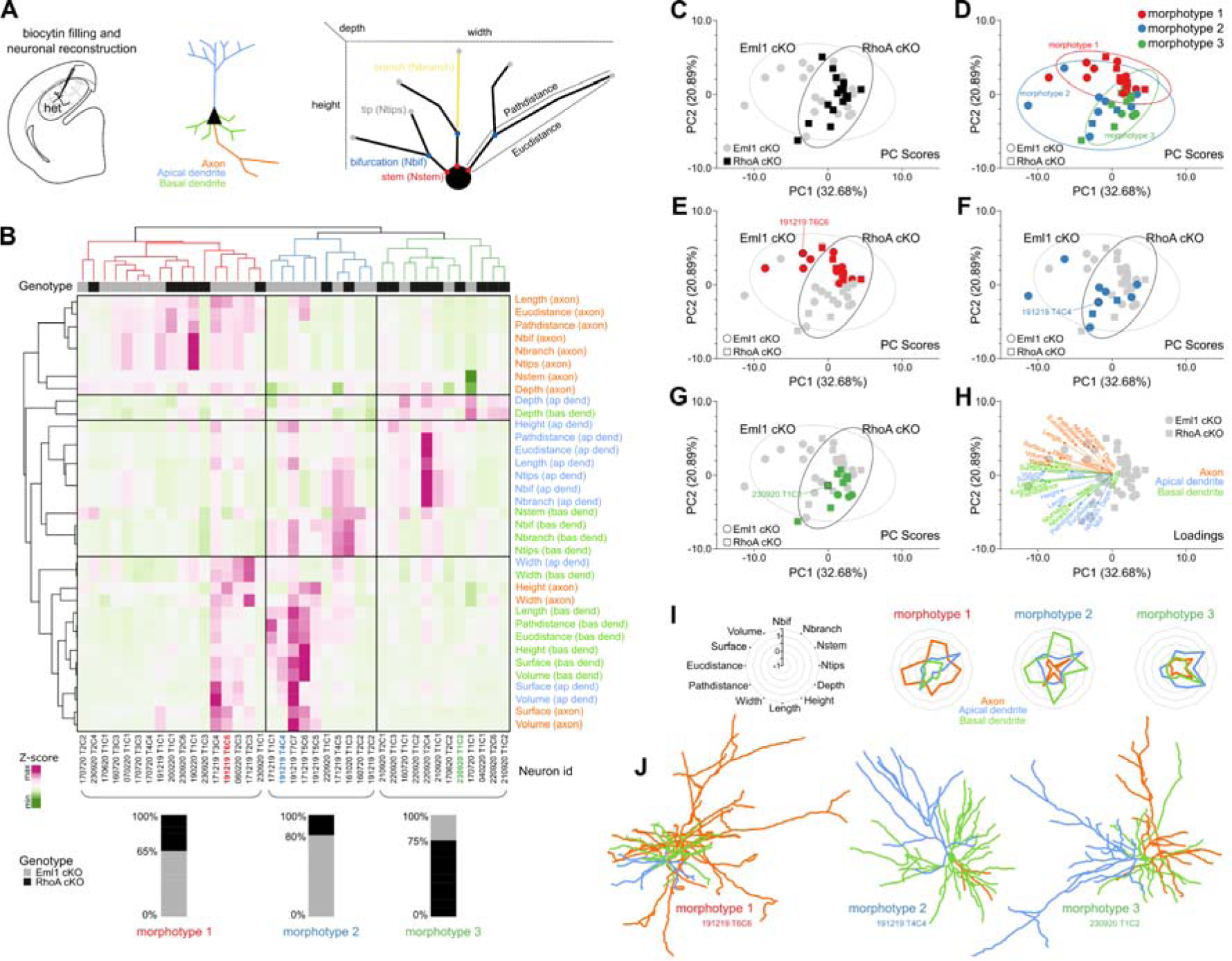
Morphological diversity in PVNH-like and SBH-like heterotopia. **(A)** Schematic views of the experimental design (left), neuronal compartments (middle) and morphological parameters extracted from neuronal reconstructions (right). (**B**) Cluster analysis dendrograms and heatmap of 39 heterotopia neurons from RhoA and Eml1 cKO mice parsed into 3 morphotypes. The genotype composition for each morphotype is shown below. (**C-H**) Principal component analysis of the same 39 heterotopia neurons from RhoA and Eml1 cKO mice with highlighted genotypes (C) and morphotypes (D-G). The loadings plot for all morphological parameters is shown in H. (**I**) Radar charts illustrating the median values for axons, apical and basal dendrite parameters for each morphotypes. (**J**) Three examples for each morphotype are illustrated, corresponding to the reconstructed neurons identified in B, E-G.

### Neurons in PVNH-like and SBH-like heterotopia have distinct electrophysiological properties

To functionally characterize heterotopia neurons, we performed whole-cell patch clamp recordings and compared the intrinsic physiological properties in the two heterotopia subtypes (Fig 3A-I). Depolarizing current injections evoked action potentials (AP) of similar amplitude in heterotopia neurons of Eml1 and RhoA cKO mice (Fig 3B, G). While heterotopia neurons in Eml1 cKO mice had a lower rheobase (Fig 3D) and a lower AP threshold (Fig 3E), their firing also showed a strong adaptation to the sustained injection of depolarizing currents (Fig 3B, C) such that heterotopia neurons from RhoA cKO mice ultimately responded with significantly more action potentials to a wide range of depolarizations than heterotopia neurons from Eml1 cKO mice (Fig 3B, C, F). Hyperpolarizing voltage pulses to evoke the hyperpolarization-activated cation current (Ih) revealed a non-significant increase of Ih current density in RhoA cKO heterotopia neurons compared to Eml1 cKO, especially at the more hyperpolarized potentials (Fig 3H, I). Other parameters, resting membrane potential (Fig 3K) and membrane capacitance (Fig 3L) were similar in the two heterotopia subtypes, whereas membrane resistance tended to be lower in RhoA cKO heterotopia neurons (Fig 3M). Overall, these results are consistent with the fact that heterotopia neurons of RhoA cKO mice have a higher excitability than those of Eml1 cKO mice.

**Figure 3.**
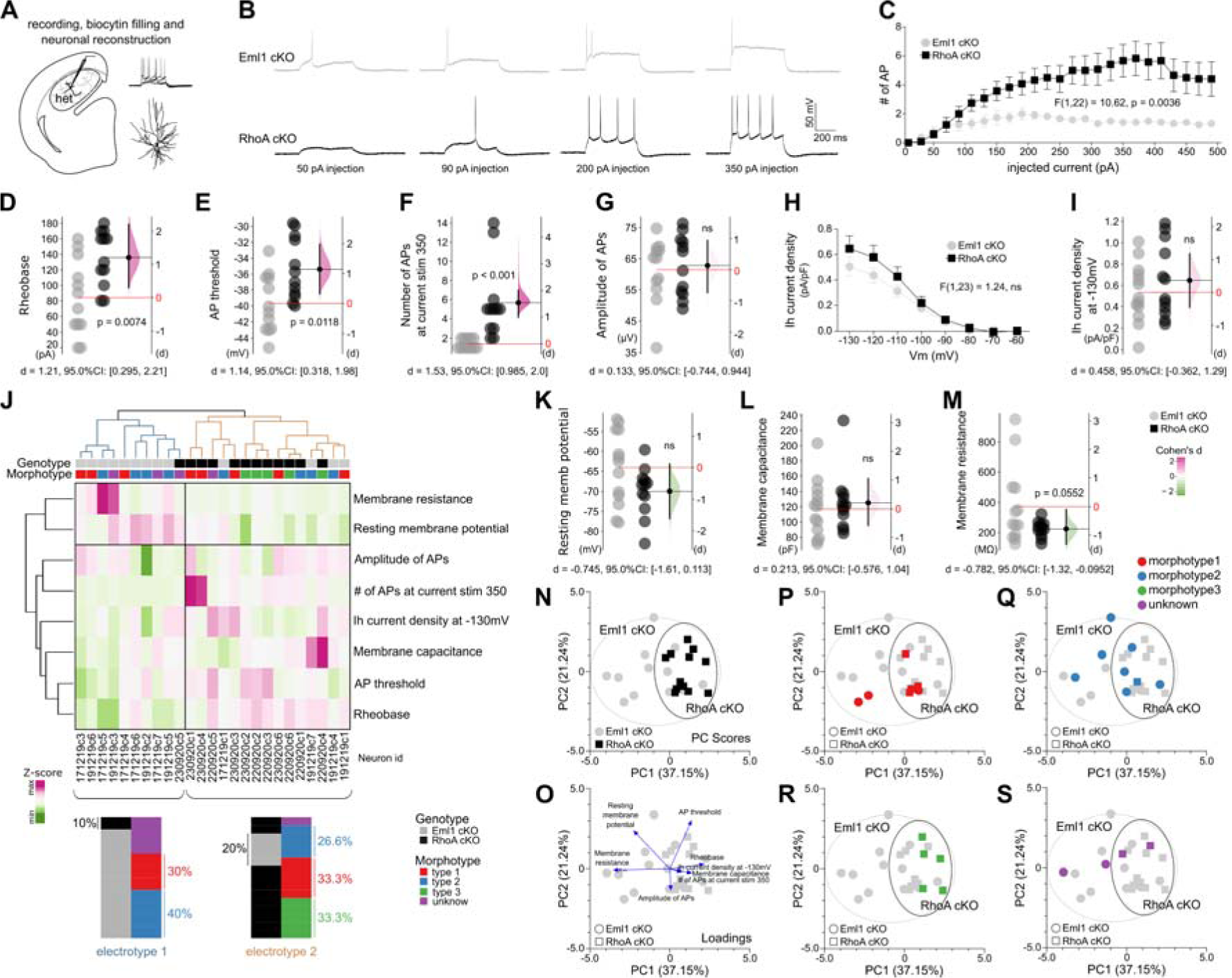
Physiological diversity in PVNH-like and SBH-like heterotopia. (**A**) Schematic view of the experimental design. (**B**) Traces of action potential firing in response to depolarizing current injections in Eml1 (top) compared to RhoA cKO (bottom) heterotopia neurons. (**C**) Input-output curves illustrating the mean number of action potential in response to depolarizing current injections in Eml1 (gray circles) compared to RhoA cKO (black squares) heterotopia neurons. Two-way repeated measure ANOVA, N=12 neurons per GMH model. (**D-G, I, K-M**) Gardner-Altman estimation plots illustrating the mean difference between Eml1-cKO and RhoA-cKO heterotopia neurons for each electrophysiological parameters, rheobase (D), action potential threshold (E), number of APs at current stimulation 350 pA (F), amplitude of APs (G), Ih current density at -130mV (I), resting membrane potential (K), membrane capacitance (L) and membrane resistance (M). Individual values for both groups are plotted on the left axes, with gray circles for Eml1 and with black circles for RhoA cKO heterotopia neurons. On the right axes, mean differences (expressed as Cohen’s d) are plotted as bootstrap sampling distributions (shaded areas). Black dots with vertical error bars show mean difference and 95%CI. Two-sided permutation t-test, N=12 or 13 neurons per GMH model (**H**) Current-voltage relationship of lh current in Eml1 (gray circles) compared to RhoA cKO (black squares) heterotopia neurons. Two-way repeated measure ANOVA, N=12 or 13 neurons per GMH model. (**J**) Cluster analysis dendrograms and heatmap of 25 heterotopia neurons from RhoA and Eml1 cKO mice parsed into 2 electrotypes. The genotype and morphotype composition for each electrotype is shown below. (**N-S**) Principal component analysis of the same 25 heterotopia neurons from RhoA and Eml1 cKO mice with highlighted genotypes (N) and morphotypes (P-S). The loadings plot for all physiological parameters is shown in O.

To assess in an unbiased fashion whether the electrophysiological properties of neurons of both heterotopica subtypes clustered in distinct ‘electrotypes’, and whether these corresponded to the morphotypes identified above, we performed hierarchical clustering of the physiological parameters (Fig 3J). We found that heterotopia neurons clustered into 2 electrotypes that closely matched their genotype, with 90% of electrotype 1 neurons being from Eml1 cKO mice and 80% of electrotype 2 neurons being from RhoA cKO mice. We confirmed these observations with principal component analysis (Fig 3N-S). From the recorded heterotopia neurons with recovered neuronal morphologies, morphotype 3 was the only morphotype found selectively associated with electrotype 2, while the two other morphotypes could be identified across the two electrotypes (Fig 3J, P-R). This finding was expected considering that each electrotype referred to a single genotype principally and because morphotype 3 was enriched in RhoA cKO mice, while morphotypes 1 and 2 were enriched in Eml1 cKO mice (see Fig 2B). Overall, these results indicate that neurons of different heterotopia subtypes have distinct electrophysiological properties, although they may share common aberrant neuronal morphologies.

To assess whether these sets of electrophysiological properties in heterotopia neurons were globally different from those previously described in cortical neurons,^41^ we performed principal component analysis. This analysis (Supplementary Fig 1A, B) not only confirmed that the heterotopia neurons of both heterotopia subtypes had different electrophysiological properties from each other, but also showed that these properties, despite some similarities (Supplementary Fig 1C), were globally distinct from those of cortical neurons (Supplementary Fig 1D-H), further highlighting their abnormal nature.

### Different dynamics of epileptiform activity in brains with PVNH-like and SBH-like heterotopia

Grey matter heterotopia are commonly associated with an increased risk for epilepsy. To evaluate whether cortical slices with heterotopia have a higher propensity for seizures, we elicited epileptiform activity with the GABA_A_ receptor antagonist gabazine. We used two-photon calcium imaging to record single-cell dynamics in cortical slices from the heterotopia models and controls. Under baseline conditions without gabazine, despite a lower number of spontaneously active cells in Eml1 cKO slices (Supplementary Fig 2A), frequencies of spontaneous cell activations in the two heterotopia models were similar than those of controls (Supplementary Fig 2B). However, under gabazine, cells in the heterotopia group had greater activation frequencies than those found in control slices, consistent with a higher excitability (Supplementary Fig 2A-D).

We next concentrated on slices with heterotopia and evaluated the contribution of the heterotopia and that of the normotopic cortex to the temporal dynamics of epileptiform activity (Fig 4A). In slices from both Eml1 cKO and RhoA cKO mice, gabazine application elicited a similar increase in the number of active cells in comparison to the baseline. This increase equally concerned the cells located in the heterotopia and those located in the normotopic cortex (Fig 4B). Likewise, cell activation frequencies in the two heterotopia models were similar in both the heterotopia and the normotopic cortex (Fig 4C).

**Figure 4.**
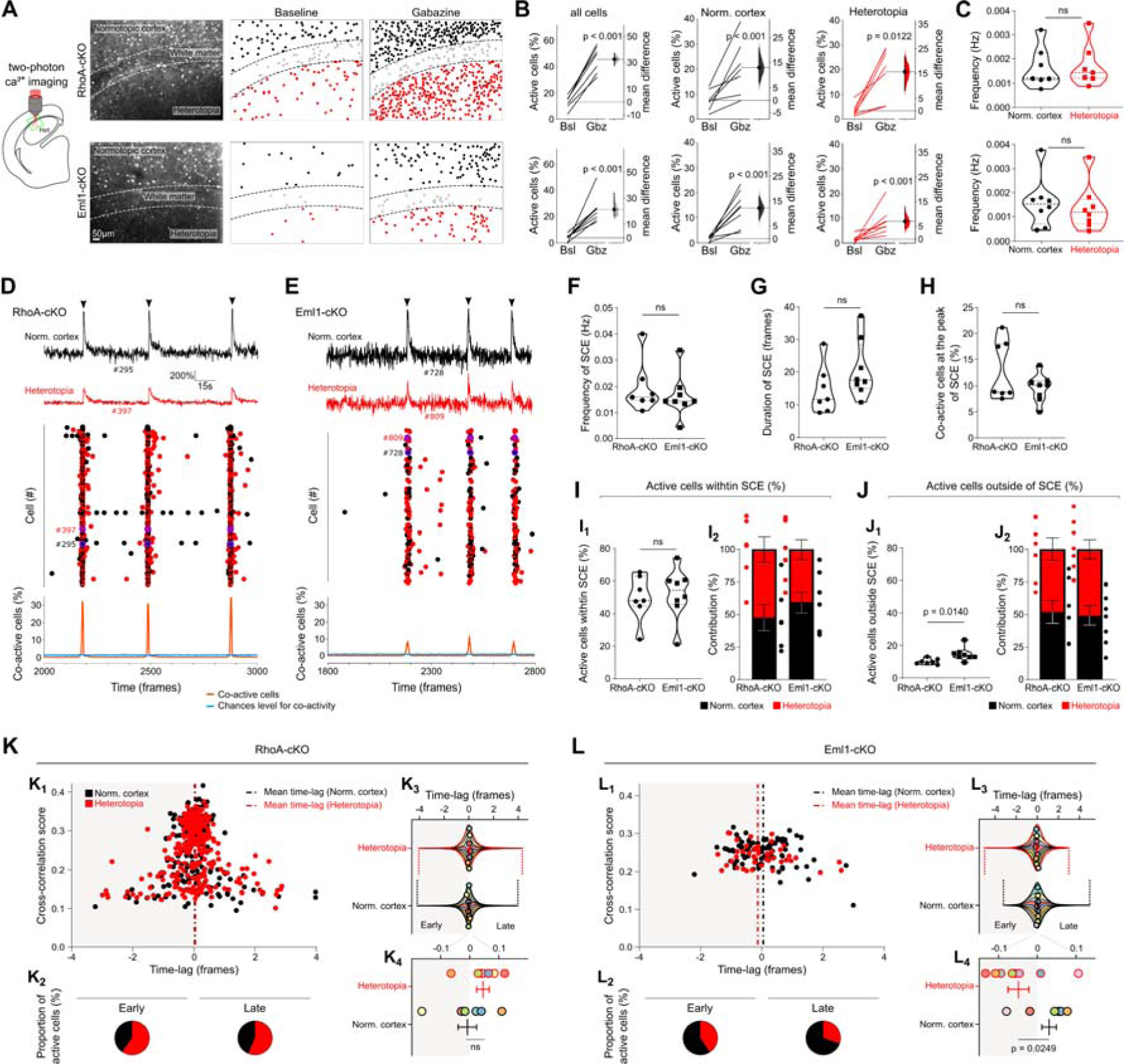
Different dynamics of epileptiform activity in brains with PVNH-like and SBH-like heterotopia. (**A**) Schematic view of the experimental design, illustrating the location of recorded areas in GMH models (left) and examples of 2-photon calcium fluorescence images (middle) in slices from RhoA (top row) and Eml1 (bottom row) cKO mice. Maps of active cells (black, gray and red dots) in baseline and under gabazine are illustrated on the right. Black dots correspond to active cells in the normotopic cortex and red dots to active cells in the heterotopia. (**B**) Gardner-Altman estimation plots illustrating the paired mean difference in the percentage of active cells between baseline and gabazine in slices from RhoA (top row) and Eml1 (bottom row) cKO mice. The left graphs illustrate all active cells, the middle graphs illustrate active cells located in the normotopic cortex and the right graphs illustrate active cells in the heterotopia. Paired sets of values measured at baseline and under gabazine are connected by a line and plotted on the left axes. On the right axes, paired mean differences are plotted as bootstrap sampling distributions (shaded areas). Black dots with vertical error bars show mean difference and 95%CI. Two-sided permutation t-test, N=7 or 8 slices per GMH model. (**C**) Violin plots illustrating the frequency of active cells in the normotopic cortex (black) and heterotopia (red) under gabazine condition in slices from RhoA (top row) and Eml1 (bottom row) cKO mice. Mann-Whitney test, N=7 or 8 slices per GMH model. (**D, E**) Top, fluorescence traces of calcium transients from individual neurons belonging to the normotopic cortex (black) and the heterotopia (red) in slices from RhoA (D) and Eml1 (E) cKO mice. Middle, raster plots of calcium events onset with black dots for cells in the normotopic cortex and red dots for cells in the heterotopia. Bottom, orange curves depict the percentage of co-active cells as a function of time (expressed as imaging frames). Blue curves depict chances level for co-activity, as estimated with a reshuffling method. Peaks of co-activity (or Synchronous Calcium Events, SCE) are defined as imaging frames during which the number of co-active cells exceeds chance levels. Mann-Whitney test, N=7 or 8 slices per GMH model. (**F, G**) Violin plots illustrating the frequency (F) and duration (G) of SCE under gabazine condition in slices from RhoA and Eml1 cKO mice. (**H, I_1_, J_1_**) Violin plots illustrating the percentage of co-active cells at the peak of SCE (H), and the percentage of active cells during (I_1_) and outside SCE (J_1_) in slices from RhoA and Eml1 cKO mice. (**I_2_, J_2_**) Bar graphs illustrating the contribution of the normotopic cortex (black) and heterotopia (red) to SCE (I_2_), and their contribution outside of SCE (J_2_) in slices from RhoA and Eml1 cKO mice. Dots show individual values, error bars show SEM. (**K_1_, L_1_**) Representative time correlation graphs illustrating, for each active cell in the normotopic cortex (black dots) and in the heterotopia (red dots), the mean time of activation (expressed as imaging frames) relative to all other cells and their cross-correlation scores in a slice from RhoA (K_1_) and from Eml1 cKO mice (L_1_). The zero-time reference corresponds to the peak of cell co-activity. Vertical dashed lines show the mean time lags for cells located in the normotopic cortex (black) and in the heterotopia (red). (**K_2_, L_2_**) Pie charts illustrating the contribution of the normotopic cortex and heterotopia to the early (top) and late phase (bottom) of SCE in the same slice as illustrated in K_1_ and L_1_. Early and late periods are defined as a function of the zero-time reference. (**K_3_, L_3_**) Violin superplots illustrating the distribution of time-lags across all experiments for cells located in the heterotopia (top) and in the normotopic cortex (bottom row) of RhoA (K_3_) and Eml1 cKO slices (L_3_). Colored stripes show the normalized density estimates for all time-lags of each slice, and outlines show the overall density estimates for all slices. Vertical dashed lines mark the left and right tails of the distributions. The superimposed open circles show mean values for each slice, and the vertical lines show mean and SEM for all slices; they are shown at an expanded timescale in **K_4_** and **L_4_**. Mann-Whitney test, N=7 or 8 slices per GMH model.

We had a closer look at the temporal dynamics of cell activations, and observed that cells were not active randomly, but instead participated in synchronous network events during which the numbers of co-active cells greatly exceeded chance levels (Fig 4D-E and methods). This is not unexpected given that recurrent synchronizations of neuronal activity are hallmarks of epileptiform activity. Frequencies of these network events were similar in the two heterotopia models (Fig 4F), whereas their durations tended to be higher in Eml1 than in RhoA cKO slices, although not reaching significance (Fig 4G). At the peak of maximum cell co-activation, we observed a tendency for greater percentages of co-active cells in RhoA cKO slices (Fig 4D, H), although similar percentages of active cells were detected within synchronous events in both heterotopia models on average (Fig 4I1). Of note, we observed significantly higher numbers of cell activations outside synchronous events in slices from Eml1 cKO mice (Fig 4J1).

We next examined the contribution of the heterotopia and that of the normotopic cortex to the dynamics of synchronous network events. On average, cells participating in synchronous events were found to equally belong to the heterotopia and normotopic cortex, both in RhoA and in Eml1 cKO slices (Fig 4I2). This suggests that in the two models, both the heterotopia and normotopic cortex contributed to epileptiform activity. To closely examine temporal sequences of cell activations during synchronous events, we calculated for each cell the average correlation score and activation time-lag relative to all other cells, as described in Marissal *et al.*^33^ We did this for cells belonging to both the heterotopia and the normotopic cortex, in the two heterotopia models, and generated time-lag correlation graphs (Fig 4K1, L1). On these graphs, the left part comprised cells recruited during the build-up of network events, while the right part comprised cells activated later on. Among these populations of early and late cells, we examined the proportion of cells belonging to the heterotopia and to the normotopic cortex. In slices from RhoA cKO mice, the proportions of heterotopia cells were higher than that of the normotopic cortex, in both populations of early and late cells (Fig 4K2). An opposite situation was observed in slices from Eml1 cKO, in which we found greater percentages of cells from the normotopic cortex (Fig 4L2). We next compared the distribution of cell activation time-lags for cells belonging to the heterotopia and to the normotopic cortex, in both models. The distributions of time-lags in RhoA cKO slices were broadly similar for cells belonging to the heterotopia and to the normotopic cortex, suggesting that neither population was recruited earlier than the other during the build-up of network events (Fig 4K3). In Eml1 cKO slices, the situation was different: while the distribution of cell activation time-lags for cells belonging to the heterotopia had a longer tail for the most negative time-lags, the time-lag distribution for cells belonging to the normotopic cortex had a longer tail for the most positive time-lags. This observation suggests that cells belonging to the heterotopia were recruited earlier than those belonging to the normotopic cortex in Eml1 cKO slices (Fig 4L3). We confirmed these results by comparing the mean activation time-lags for cells belonging to the heterotopia and to the normotopic cortex in the two models (Fig 4K3, 4K4, 4L3, 4L4). We found significantly more negative activation time-lags for heterotopia cells in Eml1 cKO slices, suggesting an earlier recruitment than for cells belonging to the normotopic cortex (Fig 4L3, 4L4). In RhoA cKO slices, we observed a tendency for a later recruitment of heterotopia cells, while not reaching significance (Fig 4K3, 4K4).

Taken together, these data indicate that both the heterotopia and the normotopic cortex contribute to epileptiform activity in the two models, but with distinct recruitment dynamics depending on the heterotopia subtypes.

### Input connectivity and axodendritic morphologies of neurons in PVNH-like and SBH-like heterotopia

To investigate the input connectivity of heterotopia neurons, we mapped functional excitatory projections to heterotopia neurons in slices from Eml1 and RhoA cKO mice using laser scanning photo-stimulation (LSPS) with glutamate uncaging. Photo-stimulations were performed at the 256 sites of a grid covering the heterotopia and the normotopic cortex to excite cells belonging to both fields successively, while synaptic responses evoked in heterotopia neurons were recorded in whole cell patch-clamp mode (Fig 5A). In slices from both Eml1 and RhoA cKO mice, we observed that the majority of inputs received by heterotopia neurons originated from the heterotopia while a minority originated from the normotopic cortex (Fig 5B, C). Because functional connectivity requires an overlap of the dendritic and axonal arbors of the connected neurons, we filled heterotopia neurons with biocytin to reconstruct and analyze their axodendritic morphologies. Consistent with the fact that the majority of inputs originated from the heterotopia, axons of heterotopia neurons mostly arborized within the heterotopia boundaries in both in Eml1 and RhoA cKO mice (Fig 5D). These observations suggested that most inputs of heterotopia neurons were from adjacent neurons in the heterotopia. However, some axons could be traced outside the heteropia limits, coursing the overlying white matter and cortex (an example is shown in Supplementary Fig 3A, together with a normotopic cortical neuron in Supplementary Fig 3B). Similarly, we observed that dendritic arbors were constrained within the heterotopia boundaries in both in Eml1 and RhoA cKO mice (Fig 5E). On average, the densities of both axonal and dendritic arbors appeared greater in heterotopia neurons from RhoA cKO mice, as compared to those of Eml1 cKO mice (Fig 5D, E).

**Figure 5.**
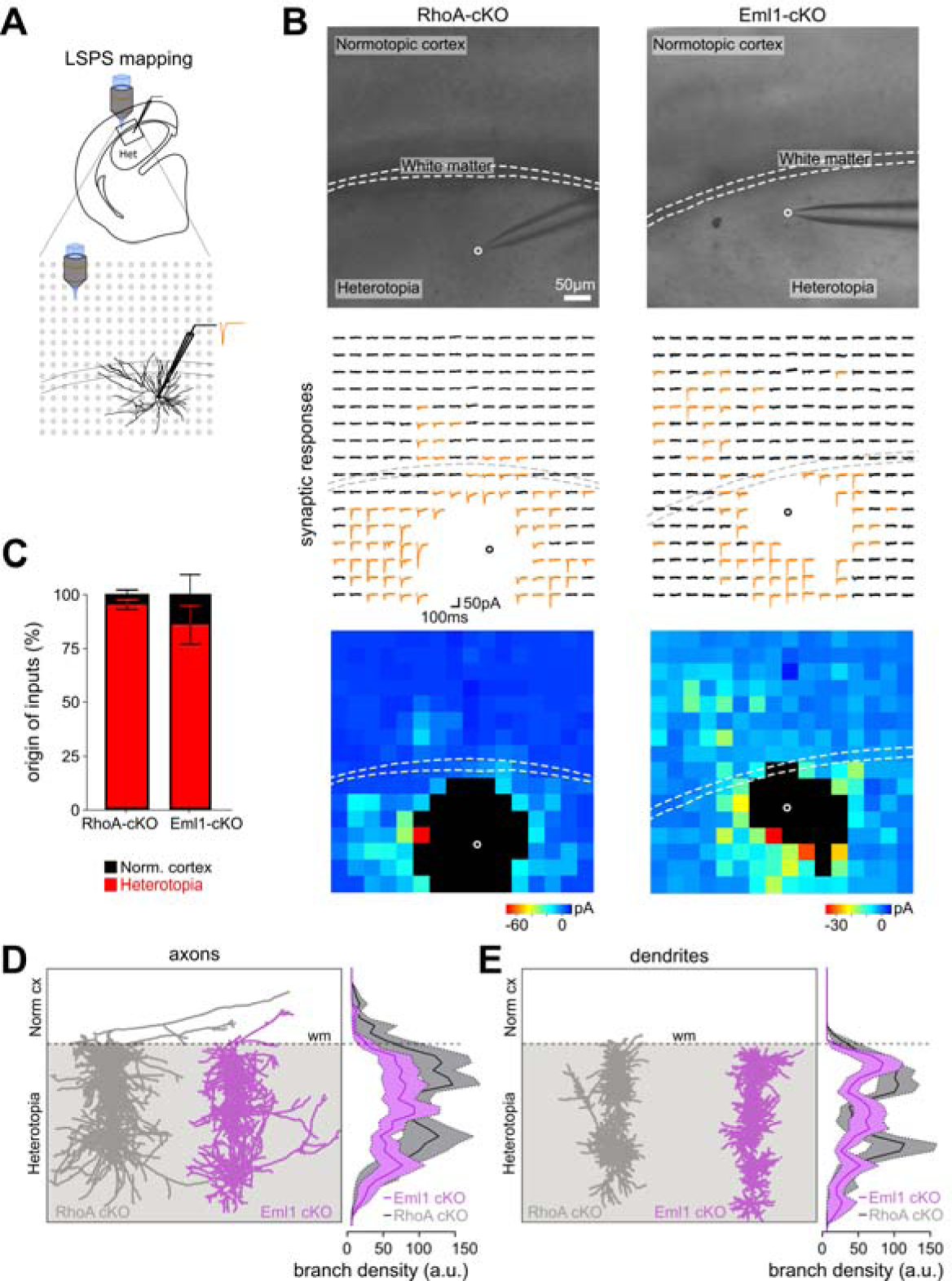
Input connectivity and axodendritic morphologies of neurons in PVNH-like and SBH-like heterotopia. (**A**) Schematic view of the experimental design for LSPS mapping, illustrating the recorded area and the location of glutamate-uncaging sites (gray dots). (**B**) Top, microphotographs showing the stimulated areas and the position of the recorded cell (open circle) in a slice from RhoA (left) and Eml1 cKO mouse (right). Middle, traces of responses of heterotopia neurons evoked by the uncaging of glutamate during LSPS. Bottom, representative input maps reporting in x and y the position of each stimulation and on a colorscale illustrating the amplitude of synaptic responses. (**C**) Bar graph illustrating the origin of synaptic inputs received by heterotopia neurons in slices from RhoA and Eml1 cKO mice. Error bars show SEM. (**D, E**) Left, reconstructed axonal (D) and dendritic arbors (E) of 17 heterotopia neurons from RhoA cKO mice (dark gray) and 22 heterotopia neurons from Eml1 cKO mice (purple). Shaded areas and dashed lines mark heterotopia boundaries. All reconstructed neurons are superimposed and projected onto a single plane. Right, line graphs illustrating the depth profile of axonal (D) and dendritic (E) branch density in heterotopia neurons from RhoA (dark gray) and Eml1 cKO mice (purple). Shaded error bands show SEM. Dash lines indicate the heterotopia border.

Taken together, these results show that the patterns of inputs received by heterotopia neurons in the two heterotopia subtypes mostly overlap with the heterotopia boundaries. This suggests that recurrent connections predominate between heterotopia neurons in both subtypes, with only a few connections received from and sent to the overlying cortex.

## Discussion

The pathophysiological basis of altered cortical function and epilepsy in GMH is poorly understood. In addition, it is unclear whether different types of GMH differ in terms of pathological consequences or, on the contrary, share common altered mechanisms. Here, we studied two robust preclinical models of SBH and PVNH, and performed a systematic comparative assessment of the physiological and morphological diversity of heterotopia neurons, as well as the dynamics of epileptiform activity and input connectivity. We uncovered a complex set of developmental changes, impacting on both the morphological and physiological properties of heterotopia neurons. Interestingly, some properties were found altered in both GMH subtypes, while others were specific to one GMH subtype. These morpho-electric abnormalities, together with altered connectivity, are likely to contribute to the different dynamics of epileptiform activity we observed between heterotopia subtypes, especially regarding the origin of onset of epileptiform events.

To our knowledge, the morpho-electric diversity of heterotopia neurons has never been investigated and compared between different types of GMH, either in resected human tissue or in murine models of GMH. Fragmented histopathological information is available in rare human surgical cases, and almost exclusively for PVNH. Analyses of the heterotopia cell type composition in human brain tissue from both children^42^ and adults^43–45^ showed that SBH^44^ and PVNH^42–46^ are composed of different types of pyramidal cells and interneurons similar to those seen in the overlying cortex, but with altered orientations and morphologies, or signs of morphological immaturity.^42^ With the exception of some PVNH cases showing signs of rudimentary laminar organization in small subcortical nodules^45^ reminiscent of chemically-induced heterotopia in rodents,^47^ no clear organization could be defined in SBH cases.^48,49^ In surgical specimens of both SBH^50^ and PVNH,^43,51,52^ pyramidal neurons whose apical dendrites can be distinguished with MAP2 immunostaining showed clearly abnormal orientations. In rodents, Golgi-stained and biocytin-filled heterotopia neurons in both SBH-like^14,53,54^ and PVNH-like^55–57^ models also showed abnormal orientations and altered morphologies similar to those found here. However, morphological similarities or differences between neurons of different GMH subtypes, as determined in our study, have not been investigated.

Due to limited access to human tissue with GMH, the electrophysiological properties of heterotopia neurons have been only studied in rodent models. Most studies have used lesional models generated using prenatal irradiation or chemicals that induce a wide range of cortical defects, including PVNH and other types of GMH such as intrahippocampal heterotopia.^58^ Targeted genetic models to characterize human mutations causing MCDs have been also generated and studied.^59^ Overall, heterotopia neurons in all models displayed variable degrees of excitability changes. Firing properties in response to depolarizing current injections have been examined both in SBH and PVNH rodent models. These studies have shown that heterotopia neurons are often heterogeneous in terms of firing patterns, with a subset of them exhibiting low-firing patterns suggestive of physiological immaturity in a genetic SBH-like model^60^ or excessive bursting patterns in a chemically-induced PVNH-like model.^55^ However, probably due to the insufficient number of recorded neurons (i.e. 3 neurons), this excessive firing behavior was not found to be associated with any specific morphological feature.^55^ Other studies reported unaltered firing properties compared with control neurons, both in a genetic SBH-like model^53^ and in an irradiation-induced PVNH-like model.^61^ However, these unaltered patterns were found to be associated with a drastic reduction of inhibitory synaptic activity, in both genetic SBH-like^53,54,60^ and irradiation-induced PVNH-like models.^61^

Collectively, these studies have clearly suggested that specific morpho-electric features may exist in different types of GMH. However, they have been carried out in models generated by different methods, either by lesions or genetic manipulation, and sometimes in different species, making a systematic comparative assessment impossible. Here, taking advantage of pertinent genetic models we have addressed this limitation by comparing in a systematic way the physiological and morphological diversity of heterotopia neurons in two models of PNVH and SBH, and have confirmed the existence of specific morpho-electric features in different types of GMH. Although this remains to be confirmed in human GMH brain tissue, a recent study in type II focal cortical dysplasia (FCD) brain tissue suggested that dysmorphic neurons belonging to the upper and lower layers show distinct morphological features,^62^ further supporting the usefulness of systematic comparative approaches in GMH and other MCDs, as we have done here.

The contribution of the heterotopia within epileptogenic networks involved in initiating seizures is unclear in patients with GMH. In PVNH patients, exploration of epileptogenic networks using stereo-electroencephalography (SEEG)^63–67^ has revealed close relationships between the heterotopia and the surrounding cortex, with epileptogenic networks encompassing both structures. Despite some degree of heterogeneity between cases, these SEEG studies in PVNH patients have shown that a large number of seizure onsets involved the normotopic cortex alone or in association with the heterotopia, while a minority of onsets involved the heterotopia alone.^43,63,66,68^ Due to the relatively small numbers of SBH cases compared to PVNH cases, and poorer outcome of surgical treatment in SBH,^69^ there are fewer reports of SEEG investigation or depth recordings in SBH patients. Similar to what has been reported for PVNH, there is heterogeneity of patients in SBH cases, with simultaneous seizure onsets in both the heterotopia and normotopic cortex^50^ or independent seizure onsets in both the heterotopia and normotopic cortex.^69,70^ Seizure onsets in the cortex and subsequent propagation to the heterotopia^71^ or no epileptiform activity from the heterotopia^69^ have also been reported.

Overall, patient reports in both PVNH and SBH suggested that the heterotopia is integrated into the same epileptogenic network as the overlying normotopic cortex and that both structures participate in epileptiform activity, as supported by observations from rodent models of PVNH and SBH.^60,72,73^ However, in both PVNH and SBH cases, the structure involved in initiating seizures varies from patient to patient, and from model to model. Such patient heterogeneity is not unexpected given the variable size, numbers and location of heterotopia, as well as different genetic causes, which are likely to contribute to making epileptogenic networks unique to each patient. In the two genetic rodent models studied here, interindividual differences between animals of the same model are minimal, and greater differences are found between models, particularly with respect to the anatomical characteristics of heterotopia and the dynamics of epileptiform activity. This is consistent with the idea that the reported patient heterogeneity would be related to the high genetic heterogeneity underlying GMH, with 146 genes and chromosomal loci identified in a recent literature review.^6^ However, future work aimed at systematically comparing different genetic rodent models of either SBH or PVNH is needed to confirm this experimentally.

Human and rodent studies have shown that epileptogenic networks in PVNH and SBH involve both the heterotopia and the surrounding cortex, thus suggesting a high degree of interrelationship between these two structures. Histopathological evidences of axonal fibers within and nearby the heterotopia have been obtained from resected human tissue using DiI tracings both in PVNH^42,51^ and SBH cases.^74^ Luxol fast blue staining for myelin revealed the presence of single or bundles of myelinated axons in PVNH,^43,51,52^ that were also visible in DiI tracings in SBH.^74^ Similar histological observations have been made in rodent models with tract-tracing studies in a chemically-induced PVNH-like model,^75^ as well as in genetic models of SBH.^16,76^ Axons of biocytin-filled heterotopia neurons were found to arborize throughout the heterotopia and send projections into the adjacent cortex in both a chemically-induced PVNH-like model^57^ and in a genetic SBH-like model,^60^ similar to what was observed here.

Analysis of structural connectivity using brain MRI in SBH patients demonstrated the presence of fiber tracts passing through or ending within the heterotopia, and also connecting the heterotopia to the overlying cortex.^77^ Similar observations have been made in PVNH cases, with fiber tracts both within the heterotopia and projecting to the surrounding white matter and overlying cortex.^78–80^ These structural connections were demonstrated to be functional with both invasive^63,64,66,81^ and non-invasive methods^79,80,82^ in PVNH cases, and non-invasive methods in SBH cases.^83,84^ Interestingly, a greater strength of functional connectivity was observed in PVNH patients with longstanding or refractory epilepsy.^79,80^ In a chemically-induced PVNH-like model, excitatory postsynaptic potentials were elicited in heterotopia neurons in response to electrical stimulation of the adjacent cortex, suggesting that these neurons received cortical inputs.^57^ The proportion of inputs received from the cortex was however not quantified. In the present study, we mapped functional excitatory projections to heterotopia neurons in both SBH and PVNH-like malformations and observed that cortical inputs are a minority in comparison to the high proportion of heterotopia inputs. Whether these proportions change with age, epilepsy onset or duration, as reported in patients^79,80^ remains to be investigated.

We recognize that there are several limitations to our study. First, to which extent the altered morpho-electric features we uncovered in the two GMH subtypes are stable or evolve with age has to be determined. As epileptogenesis in GMHs occurs in the context of ongoing cortical development and synapse formation,^85,86^ with ion channels and neurotransmitters not yet reaching their mature patterns of composition and distribution,^87^ these properties are likely to change. Whether the developmental programs are permanently altered or whether the maturation processes are simply delayed thus requires further work. Secondly, it would be worth to investigate whether seizures modify these properties, and thus alter the morpho-electric signatures. Although we did not perform longitudinal EEG recordings, we never saw evidence of behavioral seizure manifestations up to the ages we studied here (P12-15). Longitudinal electrographic recordings aimed at identifying the first signs of network dysfunction until the first seizures appear and become mature, as we recently did in a rat model of SBH,^88^ would be crucial to relate the age-dependent progression of epilepsy to specific morpho-electric signatures, if such signatures exist. Finally, how relevant are observations made in rodent models to the human pathology? There are obvious anatomical limitations to modelling MCDs in rodents, which have an unfolded cortical surface and a much smaller proportion of their cortical volume occupied by white matter than humans.^89^ This is particularly significant for GMHs, as subtypes are defined anatomically as a function of their location in relation to the white matter and ventricles. This reduced white matter volume in mouse models makes the distinction between heterotopia that are in close apposition to the ventricles, or that are separated from them by a thin line of fibers, very subtle. This is exemplified by the RhoA cKO mouse model, originally described as a SBH model.^9^ Here, we considered this model to be PVNH-like, according to recent neuropathological definitions and consensus classifications of MCDs and GMHs,^3,4^ and in particular because its most rostral ends clearly protrude into the ventricles. Using similar criteria, we considered the Eml1 cKO mouse model to be SBH-like. Although numerous reviews discuss the critical need for and usefulness of preclinical models in epilepsy research,^90,91^ to our knowledge none, except the review by Conti and Guerrini,^59^ specifically address how MCD models should capture relevant human pathology. As both the preclinical and clinical research communities need models that have construct validity, face validity and predictive value, consensus guidelines for preclinical modelling of MCDs, particularly in relation to their aetiologies and histopathological criteria, are certainly much needed in the field.

In conclusion, through a systematic comparative assessment of two mouse models, we have uncovered a complex set of altered morpho-electric features in heterotopia neurons of two GMH subtypes. Interestingly, we have found that some properties are altered in both GMH subtypes, while others are specific to one GMH subtype and result in different dynamics of epileptiform activity. Taken together, our observations may help to outline plausible scenarios for the formation of distinct pro-epileptogenic circuits in the two GMH subtypes, combining modified excitability with altered morphologies and connectivity of heterotopia neurons. While RhoA cKO heterotopia neurons tend to receive a lower density of inputs from neighboring neurons, they are more excitable, have wider apical dendrites and axonal arbors, and are therefore likely to be involved in stronger recurrent connections with other heterotopia neurons. Accordingly, RhoA cKO slices tend to involve a greater number of cells at the peak of synchronous epileptiform events, but with no clear preferential recruitment of either the heterotopia and cortex. In contrast, Eml1 cKO heterotopia neurons, although less excitable overall, comprise two morphotypes with either wider axons or wider dendritic arbors, and receive a greater density of inputs from the overlying cortex. While Eml1 cKO slices tend to recruit fewer cells at the peak of synchronous epileptiform events, these events last longer and engage heterotopia neurons with shorter delays than those of the cortex. This suggests that epileptiform events build up at a slower pace in Eml1 cKO slices, but are preferentially driven by heterotopia neurons. Overall, our work thus supports the notion that GMH represent a complex set of disorders, associating both shared and diverging pathological consequences, and contributing to forming epileptogenic networks with specific properties. Further investigation of these properties may help to refine current GMH classification schemes by identifying morpho-electric signatures of GMH subtypes, to potentially elaborate new treatment strategies. Future research may also explore possible changes in morpho-electric signatures with age, and look for potentially different signatures at different stages before and after epilepsy onset.

## Acknowledgements

We thank Prof. Cord Herbert Brakebusch for providing SC with the original breeding pairs for establishing a RhoA cKO colony. We thank the following INMED core facilities: PE and A2 animal facilities, PBMC and inMAGIC for excellent technical support. We also thank Dr. Qingzong Tseng from the CENTURI multi-engineering platform for help with image processing and analysis.

## Supplementary figures

**Supplementary Figure 1.**
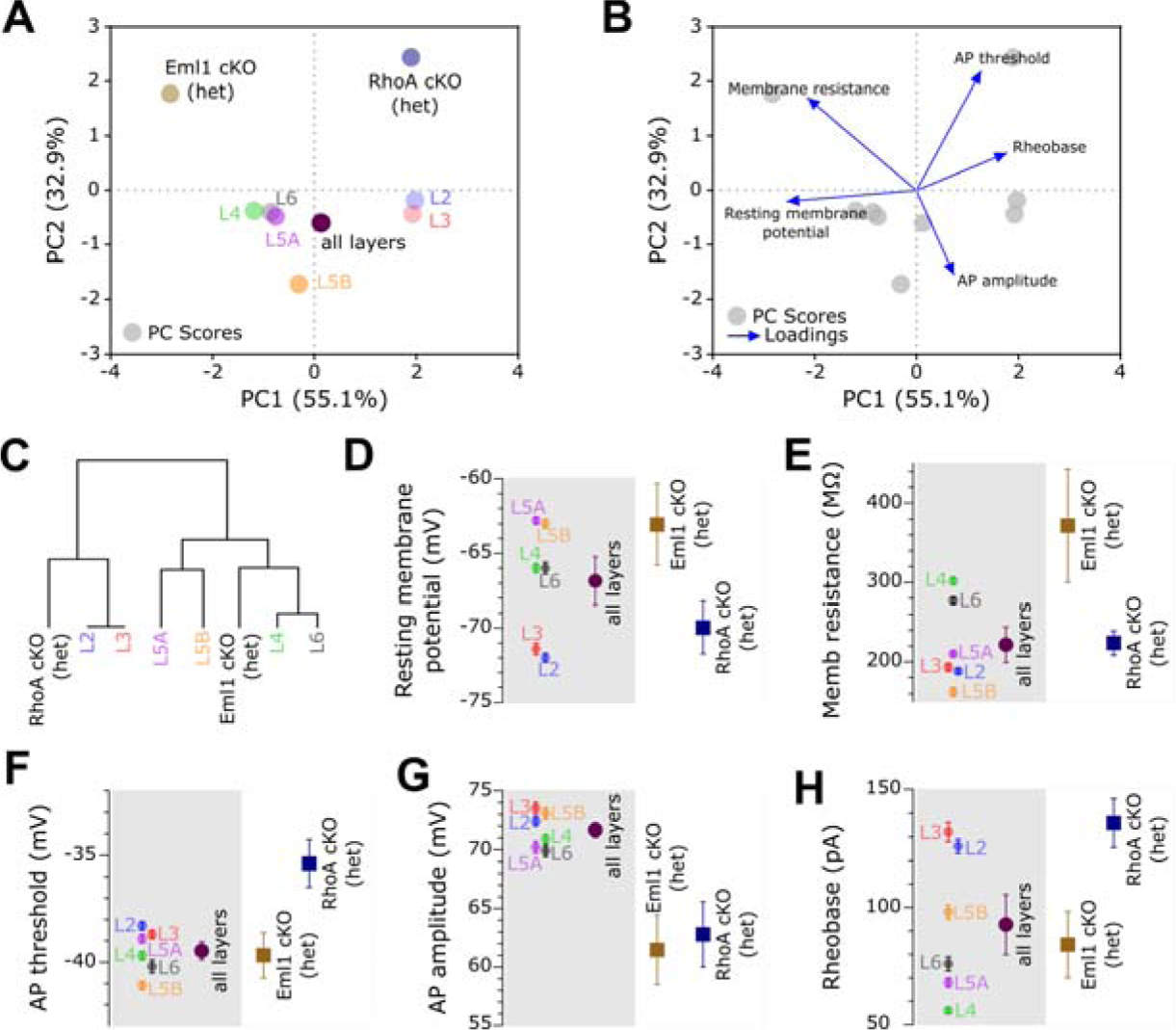
Physiological diversity of PVNH-like and SBH-like heterotopia neurons compared to that of normal cortical neurons. (**A, B**) Principal component analysis and cluster analysis dendrogram (**C**) of Eml1 and RhoA cKO heterotopia neurons and normal cortical neurons belonging to all layers, based on their previously characterized electrophysiological properties.^40^ The PC scores and loadings for all parameters are shown in B. (**D-H**) Dot plots showing the mean values for individual cortical layers or all layers combined, and that of Eml1 and RhoA cKO heterotopia, for resting membrane potential (D), membrane resistance (E), action potential threshold (F) and amplitude (G) and rheobase (H). Values for cortical neurons are taken from Lefort et al.40 Error bars show SEM.

**Supplementary Figure 2.**
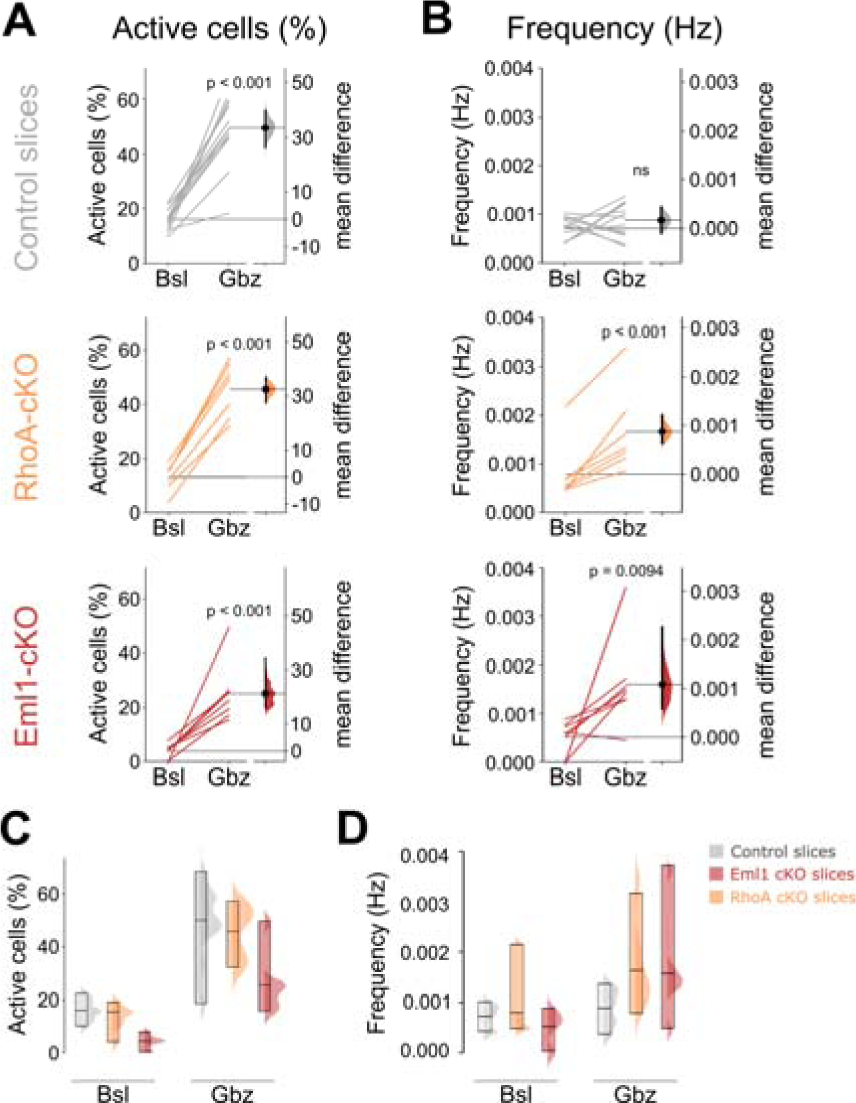
Similar number of active cells but lower cell activation frequencies in control slices. (**A, B**) Gardner-Altman estimation plots illustrating the paired mean difference in the percentage of active cells (A) and the frequency of cell activations (B) between baseline (Bsl) and gabazine (Gbz) in control slices (top row, grey) and in slices from RhoA (middle row, orange) and Eml1 (bottom row, red) cKO mice. Paired sets of values measured at baseline and under gabazine are connected by a line and plotted on the left axes. On the right axes, paired mean differences are plotted as bootstrap sampling distributions (shaded areas). Black dots with vertical error bars show mean difference and 95%CI. Two-sided permutation t-test, N=7 or 8 slices per GMH model, N=13 for control slices. (**C, D**) Bar graphs showing medians and ranges of values, superimposed with sampling distributions of the percentages of active cells (C) and cell activation frequencies (D) for control (grey), RhoA (orange) and Eml1 (red) cKO slices, in baseline and under gabazine.

**Supplementary Figure 3.**
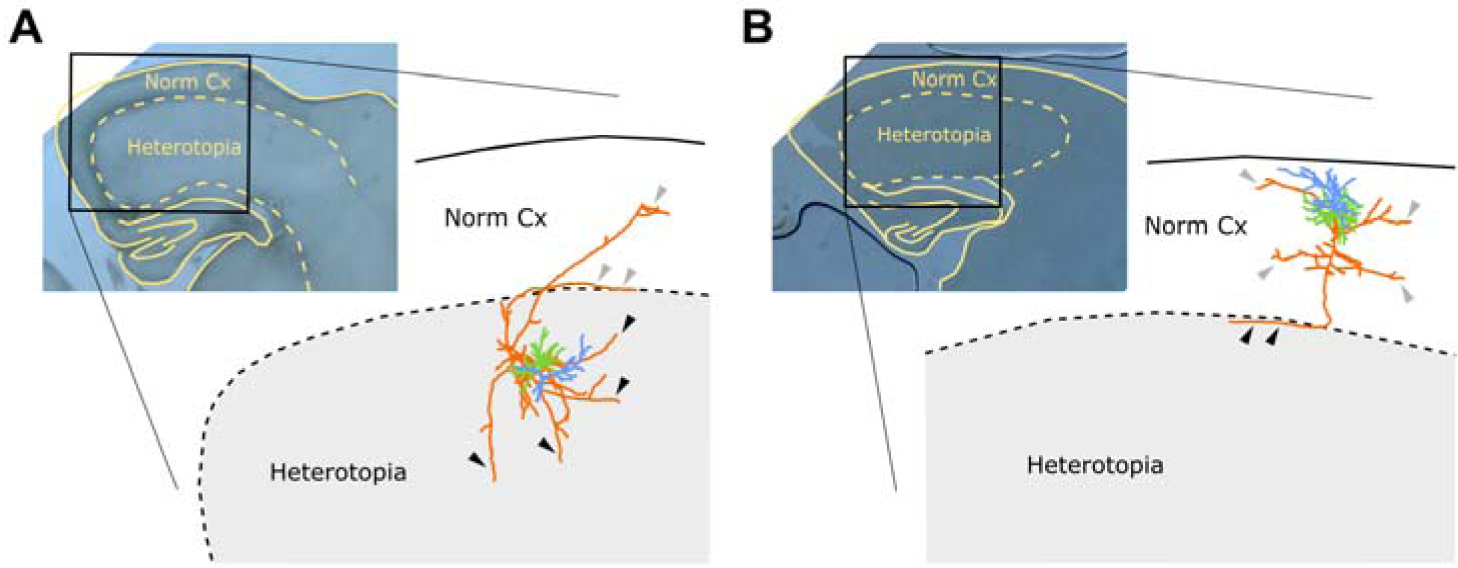
Axodendritic morphologies of heterotopia and normotopic cortical neurons. (**A**) A biocytin-filled reconstructed heterotopia neuron with its axon (orange) arborizing within the heterotopia (collaterals, black arrowheads) and sending collaterals outside the heterotopia limits (grey arrowheads), coursing the overlying white matter and cortex. (**B**) A biocytin-filled reconstructed layer 2/3 neuron of the normotopic cortex with its descending axon (orange) showing typical collateral branches in layers 2/3 and 5 (grey arrowheads), and extending down to the white matter to reach the heterotopia (black arrowheads). The two reconstructed neurons are from Eml1 cKO mice. Apical dendrites are in blue and basal dendrites in green.

